# Programmed cell death can increase the efficacy of microbial bet-hedging

**DOI:** 10.1101/059071

**Authors:** Eric Libby, William W. Driscoll, William C. Ratcliff

## Abstract

Programmed cell death (PCD) occurs in both unicellular and multicellular organisms. While PCD plays a key role in the development and maintenance of multicellular organisms, explaining why single-celled organisms would evolve to actively commit suicide has been far more challenging. Here, we explore the potential for PCD to act as an accessory to microbial bet-hedging strategies that utilize stochastic phenotype switching. We consider organisms that face unpredictable and recurring disasters, in which fitness depends on effective phenotypic diversification. We show that when reproductive opportunities are limited by carrying capacity, PCD drives population turnover, providing increased opportunities for phenotypic diversification through stochastic phenotype switching. The main cost of PCD, providing resources for growth to a PCD(-) competitor, is ameliorated by genetic assortment driven by population spatial structure. Using three dimensional agent based simulations, we explore how basic demographic factors, namely cell death and clonal reproduction, can create populations with sufficient spatial structure to favor the evolution of high PCD rates.

## Introduction

Programmed cell death (PCD) describes a genetically encoded process of cellular suicide that is often used as an umbrella term for more specific cell-death phenotypes (e.g., apop-tosis, paraptosis, autophagy, chromatolysis, etc.) [1, 2, 3, 4, 5]. Anatomists first observed PCD in the context of animal development during the 19th century [4]. Since then, a vast body of literature has established the key role of PCD in both the generation [6, 7] and maintenance of multicellular forms [1, 8]. Interestingly, PCD is also widespread among distantly related unicellular organisms [9, 10, 11, 12, 13, 14, 15, 16]. The origin and maintenance of PCD within multicellular taxa has a straightforward evolutionary explanation if the death of some cells provides a benefit to the organism as a whole. In contrast, the evolution of PCD in unicellular organisms presents a conundrum: under what conditions (and by what mechanisms) will natural selection favor organismal suicide?

Different mechanisms have been proposed to explain the existence of PCD among unicellular taxa [15]. One category of hypotheses proposes that PCD is an altruistic trait favored by kin or multilevel selection. These hypotheses propose that PCD may have evolved to remove cells weakened by deleterious mutations, pathogens, or age-accumulated damage [17, 18, 19, 20, 21, 22]. Removing such cells improves the health of other members of the population either by preventing the spread of pathogens or making more resources available to healthier cells. Another category of hypotheses considers PCD to be a pleiotropic side-effect of genes under positive selection because of their pro-survival effects [15]. This would imply that there is no direct adaptive benefit to PCD and its negative effects are simply a tolerable side-effect of a beneficial pleiotropic trait. Finally, PCD may have evolved in microbes because of its role in multicellular development. For example, PCD by a subset of a bacterial population may be necessary to provide extracellular DNA that plays a structural role in biofilm formation [23]. Unfortunately, few of these potential evolutionary explanations have been experimentally tested or mathematically modeled, and little is known about the ecological conditions necessary for their evolution.

In this paper, we propose a novel evolutionary hypothesis for the origin and maintenance of PCD in unicellular organisms: PCD serves as an accessory to microbial bet hedging. Bet-hedging traits increase fitness in unpredictable environments in two possible ways. First, they can spread risk among multiple phenotypes, each of which is well-suited to a possible future environment (diversification bet-hedging) [24, 25]. Second, they can allow organisms to pursue a generalist strategy that performs acceptably across a range of possible future environments (conservative trait bet-hedging) [26]. Of the two types of bet-hedging, most of the well-established traits act as diversification bet-hedging, but this may be because it is more conspicuous than conservative trait bet-hedging [27]. Micro-organisms typically enact diversification bet hedging strategies through stochastic phenotype switching, in which reproducing cells give rise to phenotypically distinct offspring with a low (typically 10^−1^ to 10^−5^) probability [28, 29]. Since the offspring can switch back to the original phenotype at some low probability, stochastic phenotype switching typically generates bistable populations in which a single genotype exhibits two distinct phenotypes [30, 31]. Importantly, stochastic phenotype switching requires generational turnover to create variation. Even at relatively high rates of switching (10^−3^), it still takes more than 1,000 generations for an initially uniform population to reach maximum levels of phenotypic diversity [28].

Here we examine the conditions under which PCD increases the efficacy of microbial bet-hedging by creating generational turnover, resulting in increased phenotypic diversity. We analytically examine the co-evolution of PCD and stochastic phenotype switching in an unpredictable environment in which more diversified populations have higher long-term fitness. Although population size in our model is constrained by a carrying capacity, PCD allows organisms to circumvent this limitation to reproduction. As organisms die, they create opportunities for other organisms to reproduce and diversify via stochastic phenotype switching. Thus PCD incurs both costs and benefits: some cells die, but if surviving clonemates can use spared resources to divide, then the genotype as a whole will become more diversified.

One possible downside of this strategy is that the resources made available by PCD are susceptible to exploitation by low-PCD competitors. We show that across many cycles of unpredictable environmental risk, exploitation by low-PCD competitors does not necessarily overwhelm the long-term fitness advantage gained by the more diversified high-PCD strain. Further, the cost of PCD is highly dependent on the degree of population structure and is reduced when individuals that die are more often replaced by nearby, related clonemates. More importantly, we find that the conditions required for selection to favor elevated PCD in our model are very permissive: elevated PCD can evolve in microbes with a wide range of stochastic switching frequencies, in environments with a wide range of disaster frequencies, and in populations with modest spatial structuring. We contextualize these results using a spatially explicit biofilm simulation. We find that, even if the simulation is initiated with a well-mixed population, the dynamics of occasional environmental catastrophe and range expansion creates high levels of spatial structure, which rapidly favors the evolution of high rates of PCD. These results point to new adaptive explanations for the evolution and maintenance of PCD in unicellular populations by focusing attention on the profoundly non-equilbirium nature of many microbial populations (particularly those exploiting patchily-distributed, ephemeral resources).

## Results

### Model

We consider a competition between two microbial strains *(G1* and *G2)* in an environment that experiences frequent disasters (previously described in [28]). Each strain exhibits two possible phenotypes: A and B. We denote the A and B phenotypes of the G1 genotype as *A1* and *B1* and similarly A2 and B2 correspond to the phenotypes of G2. The only meaningful difference between *A* and *B* phenotypes is their susceptibility to an environmental disaster. Disasters occur randomly and target a single phenotype for annihilation-in this way they are similar to other disasters that microbes might face such as antibiotic exposure or immune system recognition. Following a disaster, the surviving types reproduce until the total population is restored to some fixed amount, N. Phenotypic diversification occurs via stochastic phenotype switching such that each time a strain reproduces there is a probability (*p*_1_ for *G1* and *p*_2_ for *G2*) that the alternate phenotype will be produced. In between disaster events, there are rounds of PCD in which each strain loses cells according to their own characteristic PCD rate, *c*_1_ and *c*_2_. The population is restored to *N* in a way that incorporates environmental structure or relatedness, characterized by a structure parameter *r* ∊ [0,1] (shown in Figure 1). The r parameter determines the fraction of cells lost to PCD that will be replaced by the same genotype. If *r* = 0, then no cells that died via PCD will be replaced by the same genotype; if *r* = 1 then all cells will be replaced by the same genotype. The equations in Eqn. 1 describe the dynamics of a single round of PCD and repopulation. Discrete rounds of PCD and disasters continue until one or both genotypes go extinct. A genotype is said to “win” if the opposing strain goes extinct first or if at the end of a thousand rounds it makes up a higher percentage of the total population. Simulations were coded in the language python and are available in the Supplementary material.

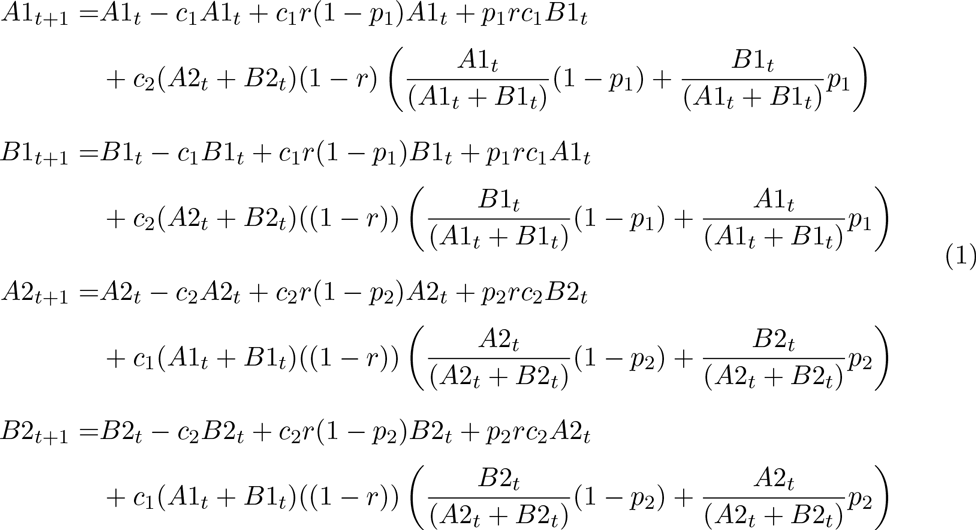

### Cost of PCD

There is a direct cost to PCD because cells are killed. If all cells were replaced by the same strain then this would remove the cost. However, if replacement does not occur with perfect assortment (i.e. *r* < 1) then some of the cells that die will be replaced by members of the competing strain. We call the total number of a strain that are replaced by a competing strain the cost of PCD. Since our model has disasters interspersed with rounds of growth and PCD, we can calculate the expected cost of PCD during the period between disasters. If one strain uses PCD and the other does not then after *t* rounds of PCD/regrowth the PCD strain will decrease by a factor shown in Eqn. 2.

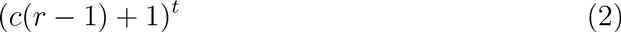

**Figure 1:**
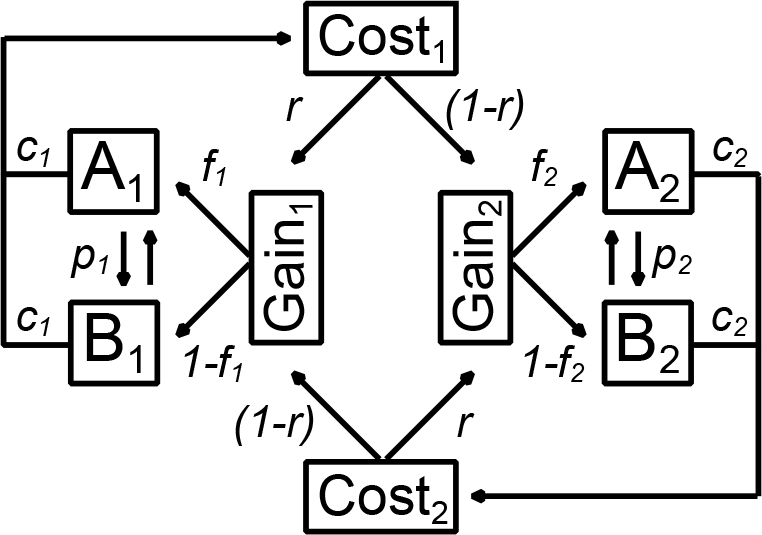
Model schematic for the dynamics of the competition Genotypes. *G1* and *G2* switch between phenotypes *A* and *B* with a probabilities *p_1_* and *p*_2_, respectively. Organisms undergo PCD with probabilities c_1_ and *c*_2_ such that *c*_1_(*A*_1_ + *B*_1_) is the expected “cost” of PCD experienced by genotype *G1*. The population then regrows back to the carrying capacity N such that each genotype repopulates a fraction of its own cells that underwent PCD determined by the value of *r*. The repopulation is partitioned among phenotypes according to their relative frequency such that 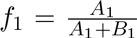 and 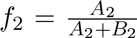. The “gain” corresponds to the number of cell reproductive events reallocated to each genotype.

This factor has three terms which determine its magnitude: *c*, *t*, and *r*. As expected, the cost of PCD is high for large values of *c*, i.e. the PCD rate. Similarly, if disasters are infrequent, i.e. *t* is large, then the cost is high because it effectively increases the total amount of PCD that occurs in between disasters. Since the *r* parameter determines how many cells that undergo PCD are replaced by the same genotype, higher values of *r* reduce the cost of PCD. When *r* = 1, Eqn. 2 is equal to 1 which means that there is no cost to PCD. A parameter that is missing from the cost factor is the switching rate, *p*. This is because the cost is absorbed by the strain as a whole without regard to how it is partitioned into *A* and *B* types.

### Benefit of PCD

While the cost of PCD is shared by all members of a genotype without regard to phenotype, the benefit of PCD is determined by diversification among phenotypes. In our model, PCD allows a strain more chances to diversify via stochastic phenotype switching by increasing the number of reproductive events between disasters. Such diversification could manifest in a more equal mix of the population among phenotypes, which can increase long-term fitness by reducing variance in fitness across disasters. Alternatively, if there are few opportunities for diversification between disasters (for example, strains with a very low rate of stochastic switching), higher diversification rates can reduce a strain’s possibility of extinction. We will focus on this second manifestation because it is more crucial to the survival of the PCD strain— failure to diversify can result in extinction.

We assume that one phenotype, say *B*, has just experienced a disaster and is no longer present in the population. Each strain only exists as the *A* phenotype with m members of the PCD strain (*A*_1_) and *N – m* of the non-PCD strain (*A*_2_). Prior to the next disaster, each strain must produce *B* phenotypes or face extinction should the next disaster target *A* phenotypes. Although there may be several rounds of PCD/regrowth in between disasters, we investigate the consequences of just one round of PCD/regrowth. We calculate the expected number of *B* types produced by each genotype assuming that the switching probability is the same, i.e. *p*_1_ = p_2_ = *p*. The number of B types produced by the PCD genotype (B1) is prC_1_m and the non-PCD genotype (*B*2) is *p*(1 — *r*)*c*_1_*m*. The difference between *B*1 and *B*2 is determined solely by the r parameter. We note that the *B*2 types produced by the non-PCD genotype only happen as a result of PCD undergone by *G*1 where replacement was not perfectly assortative (*r* < 1).

**Figure 2:**
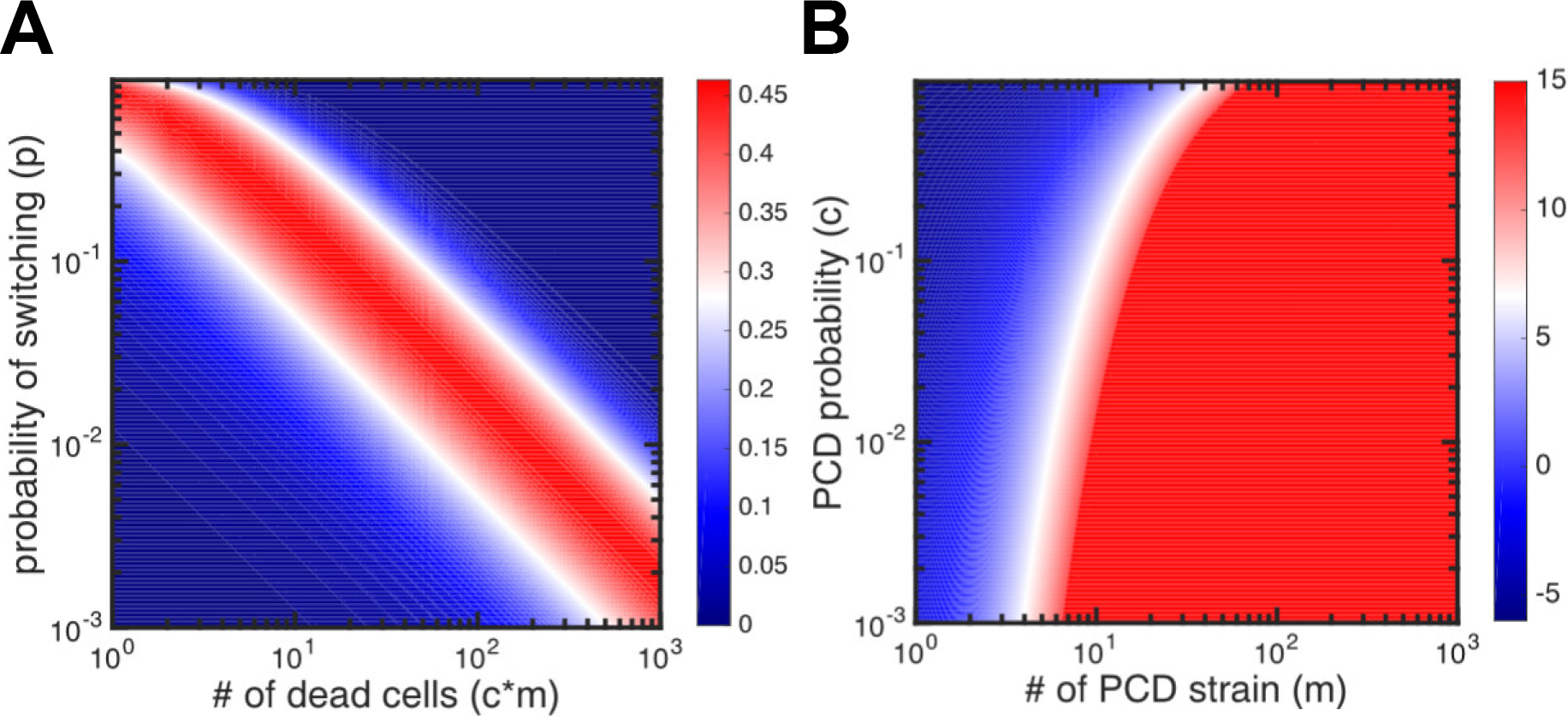
Cost and benefit of PCD. A) A contour plot of the benefit of PCD from Eq. 3 for a range of switching probabilities (*p*) and numbers of PCD cells (*c* * *m*) when *r* =.75. There is a narrow band where PCD is beneficial corresponding to *cmp* = const. Outside of this area, the switching rate is too high or too low to incur a benefit to PCD genotypes. B) A contour plot shows the *log_10_* relative benefit versus cost of PCD as expressed in Eq. 5 for a range of PCD probabilities (*c*) and number of cells (*m*) with *r* =.5 and *p* = 10^−6^. The blue area corresponds to a greater cost while red areas correspond to a greater benefit. Since the plot is transformed by *log*io the benefit is many orders of magnitude (> 10) greater when the number of cells is larger than 10.

One advantageous situation for the PCD strain would be if it produced a *B* phenotype but the non-PCD strain did not. The probability of this event is shown in Eqn. 3.

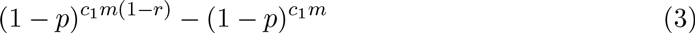

The probability increases with the structure parameter *r*, reaching a maximum at *r* = 1. It is highest at *p* = 1 − (1 − *r*)^1^*c*_1_*mr* which for *r* << 1 is approximately one expected B phenotype produced by the PCD strain (see Figure 2A). At slower switching rates and lower rates of PCD, the PCD strain is not likely to diversify. At higher switching rates and higher rates of PCD the non-PCD strain is likely to diversify along with the PCD strain, thereby reducing the relative advantage of PCD.

The probabilistic formulation of the benefit of PCD can also be compared with a probabilistic formulation of the cost. We put the cost into a similar currency by considering the probability that the PCD strain goes extinct because of PCD (shown in Eqn. 4). This requires all m cells to undergo PCD and be replaced by the competing strain.

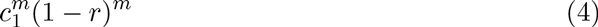

In this formulation, the PCD strain faces the cost of going extinct from PCD but has the benefit of diversifying when the competition does not. The relative probability of the benefit to the cost is shown in Eqn. 5.
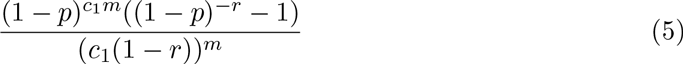

This relationship is low when *p* is very high (*p* = 1) or when *p* is very low *p* = 10^−6^ and *r* is small (*r* =.5). If we assume a low *p*, say *p* = 10^−6^, and an unbiased *r*, i.e. *r* =.5, then we can compare the ratio of probabilities for a range of values of *c* and *m* (see Figure 2B). Effectively, if *m* is larger than 10 then *the benefit is more likely than the cost by over 10 orders of magnitude.*

### Interplay between cost and benefit

While the benefit of PCD can outweigh the cost under some circumstances (Figure 2B), it is not clear how often those circumstances arise. In a competition, it could be that it is rare for both organisms to only exist in one phenotype after a cycle of a disaster and restoration to carrying capacity. To evaluate how a genotype with PCD fares in competition, we compete a PCD strain versus a non-PCD strain over a range of switching probabilities (*p*) and PCD rates (*c*). The results shown in Figure 3A reveal that the PCD strain outcompetes the non-PCD strain for intermediate switching probabilities when PCD is not too frequent, i.e. *c* <.1. For PCD rates above.1, the cost is too high to compensate for any benefit. For high switching probabilities, the benefit of PCD is diminished because it is likely that the non-PCD strain will always diversify. Similarly, if the switching probability is too low then the PCD strain cannot adequately diversify. For more structured environments, with say *r* =.9 shown in Figure 3B, the PCD strain wins over a much larger area of parameter space. This is because the higher value of *r* reduces the cost to PCD. If we fix the rate of switching to *p* =.1 then we see that as the structure parameter increases, larger values of c can be tolerated, i.e. higher PCD genotypes are successful (Figures 3C and 3D). In addition, with increasing frequency of disasters, genotypes with higher values of PCD can outcompete non-PCD genotypes— the benefit of diversification via PCD outweighs its cost.

### Cost when both strains have PCD

Until now we have considered the case in which there is only one genotype that undergoes PCD. Here, we investigate what happens when both strains exhibit PCD but at different rates, i.e. *c*_1_ > *c*_2_ > 0. Similar to the derivation of Eqn. 2, we assume that both populations undergo t rounds of PCD and regrowth. Genotype *G*1 starts with a population of *G*1_0_ and genotype *G*2 begins with *N* — *G*1_0_. We compute the number of *G*1 genotypes at time t, *G*1_t_, and find that this depends only on the PCD rates, the structure parameter *r*, and the initial amounts of each genotype (see Eqn. 6).

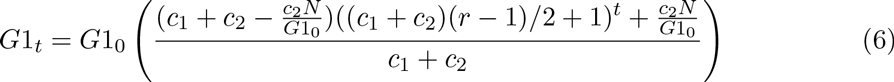

One interesting consequence of Eqn. 2 is that the higher PCD genotype, G1, can actually increase in abundance over time. This occurs because the total lost by each genotype to PCD depends on their abundance. With a fixed population size, as one genotype becomes less abundant it pays less of a cost for PCD. In Eq. 6, PCD is a cost for *G*1 if the term in parentheses is less than 1 which occurs if 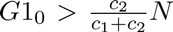. If the opposite is true then there is actually a net flux to *G*1 and it can increase in number. Without the disruption of a disaster, the genotypes approach the equilibrium shown in Eqn. 7 (assuming *c*_1_ < 1 and *r* >-1).

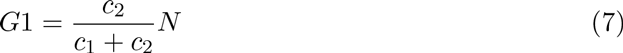

**Figure 3:**
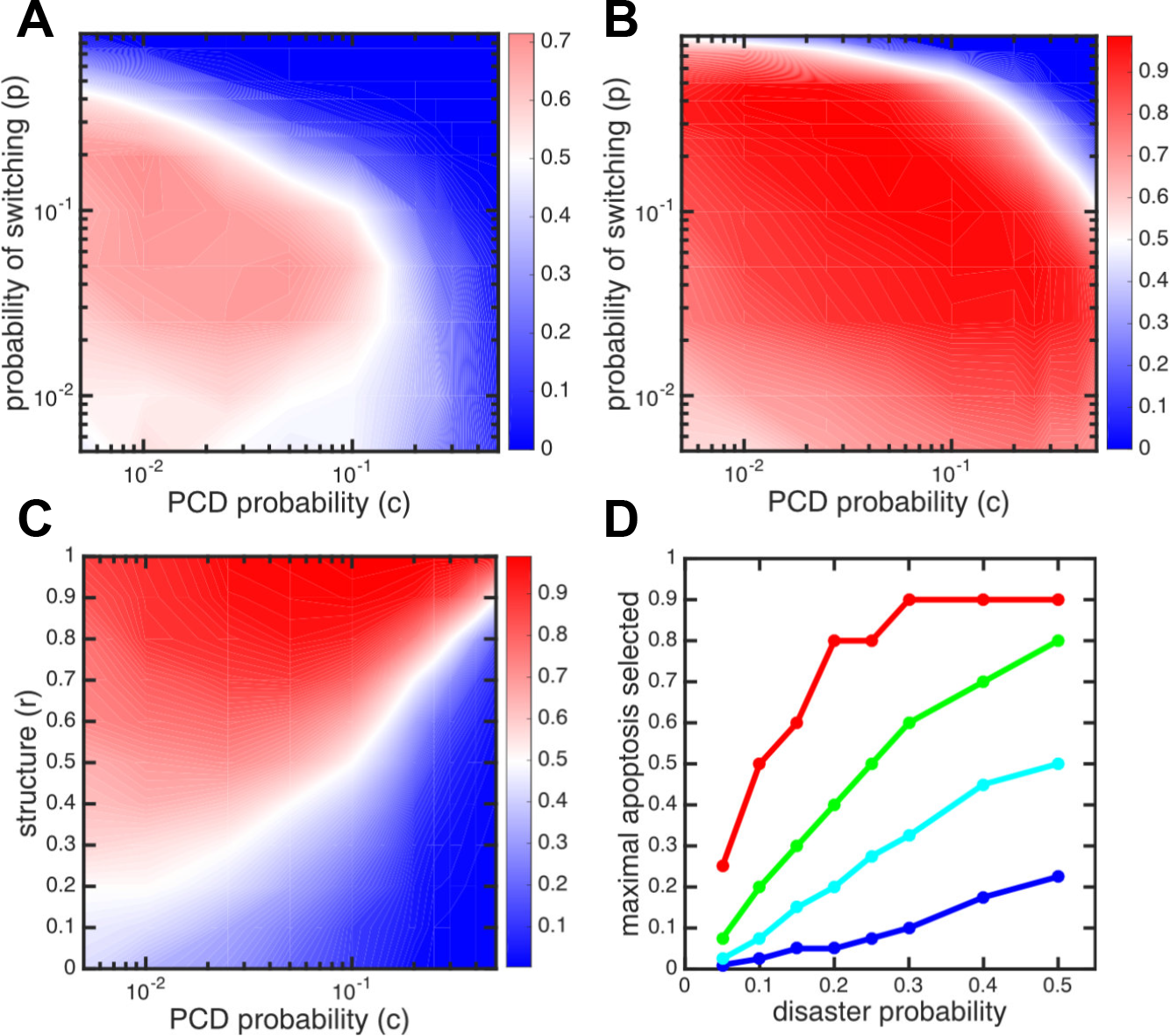
Competitions between PCD versus non-PCD genotypes. A) Contour plot shows the percentage of wins for the PCD strain versus the total number of competitions that ended in one strain winning, over a range of switching probabilities (*p*) and PCD probabilities (*c*) with *r* =.5 and disaster probability of.1. There is an intermediate range of switching probabilities and PCD probabilities where the PCD strain is more successful. The blue area corresponds to where PCD loses to non-PCD cells and the red area is where PCD wins. B) Same as A) but with *r* =.9. PCD is more successful (higher number of wins) over a larger area of parameter space. C) The success of PCD strains versus a non-PCD competitor are shown as a function of PCD rate c and the assortment parameter *r*. The disaster probability is.1 and the switch rate for both genotypes is.1. With increasing *r* PCD is more beneficial. D) The maximal amount of PCD selected is shown as a function of disaster probability for different values of r3.3 (blue), .5 (cyan), .7 (green), and .9 (red). Higher values of PCD are permitted for higher frequencies of disaster where there is greater benefit for diversification and in more structured environments where there is a lower cost to PCD.

We note that the equilibrium does not depend on the value of the structure parameter r as long as *r* > 0. Thus, as long as there is some chance that the higher PCD genotype replaces some of the population it lost to cell death, then this equilibrium will be reached. We note that in this calculation (in the absence of disasters) if *G*2 does not engage in PCD, i.e. *c*_2_ = 0, then *G*1 will eventually go extinct. If, however, both genotypes engage in PCD, then they can coexist— at least until a disaster comes.

The above calculations show the possibility of coexistence in the absence of disasters and stochasticity. We might expect that the combination of disasters and stochastic events (phenotype switching) may interfere with long-term coexistence. Figure 4 shows an example of coexistence between two PCD strains (*c*_1_ = .3 and *c*_2_ = .1) in a stochastic simulation for 10^5^ rounds. Disasters occur with a frequency of .1 which means on average we expect 10^4^ disasters. Both genotypes switch with the same probability p = .1. Despite the frequency of disasters and the different PCD rates, neither genotype goes extinct. Since the simulations are stochastic, at some point one genotype will go extinct but the waiting time until this event occurs can be long and require more than tens of thousands of disasters.

### Generation of spatial structure

Our analyses thus far have relied on a notion of spatial structure *r* that determines how much of the resources made available by PCD actually return to the genotype liberating them through programmed cell death. Although our parameter *r* is a mathematically convenient macroscopic description of population spatial structure, it is a complex variable that is both an outcome and driver of individual-level (e.g., birth/death, motility) and population-level (e.g., selection, disturbance) processes. To examine whether sufficiently high values of *r* that favor the evolution of PCD can emerge from demographic processes of cell death and local reproduction, we construct a 3D agent-based model (ABM).

**Figure 4:**
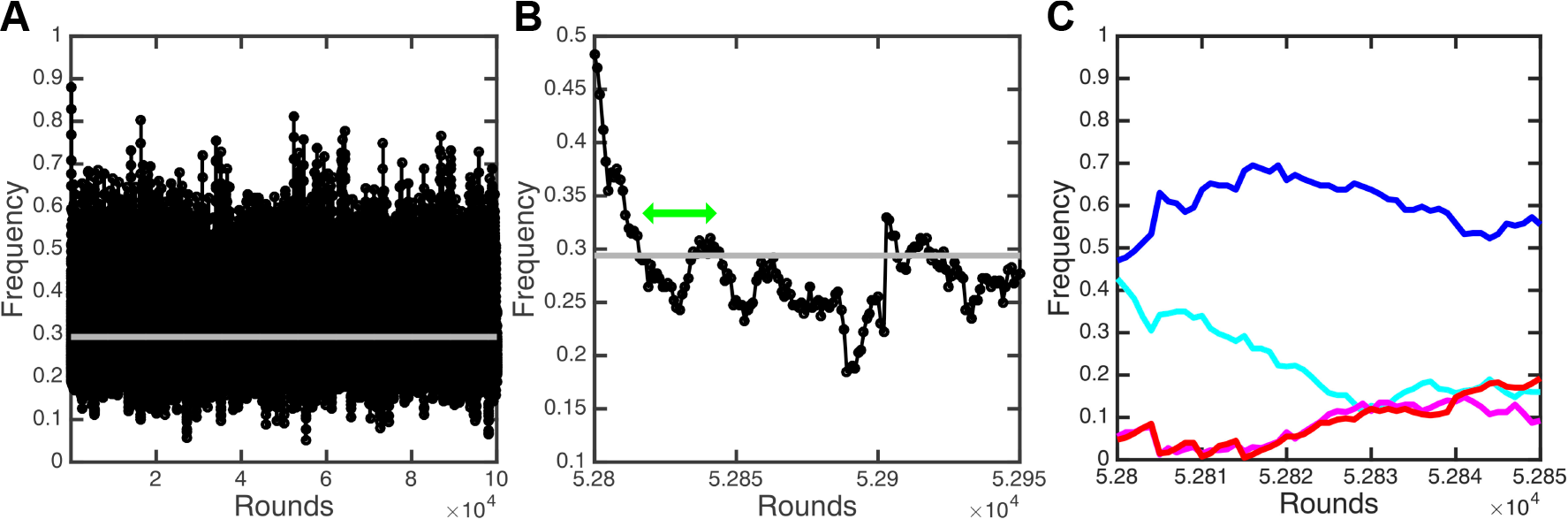
Coexistence of PCD strains. A) Two apoptotic strains with the same switching rate but different PCD rates (*c*_1_ = .3, *c*_2_ = .1) compete for 10^5^ rounds without either going extinct. The switch probability is the same for both *p* = .1 and the disaster probability is .1. The black lines show the frequency of *G*1 as it fluctuates over time, rarely getting close to extinction. The gray line shows its average frequency over the simulation. B) A highlighted segment of the competition shows that as *G*1 gets low in frequency it is bolstered up by *G*2. An indication of this event can be seen in the area denoted by the green arrow. C) The dynamics of the genotypes separated into their A and B phenotypes are shown for the area denoted by the green arrow from B). The *G*1 phenotypes A (magenta) and B (cyan) and *G*2 phenotypes A (red) and B (blue) change in frequency as a result of PCD events. *G*1 and *G*2 experience the same increase in A phenotypes (red and magenta) due to differences in PCD rates and flux between genotypes. Even though the number of B for *G*2 (blue) is greater than the B for *G*1 (cyan), the A types recover by a similar amount— this corresponds to the populations approaching the equilibrium of coexistence.

The agent-based model consists of cells distributed across patches that represent microenvironments within 3D space (see Figure 5a). Each patch can hold 10 cells at which point it reaches carrying capacity. Cells are mobile agents capable of reproduction and death. As in our mathematical model, there are four types of cells corresponding to two genotypes, each of which produce two phenotypes that are defined by their resistance/susceptibility to disasters. The main difference between genotypes is that one undergoes PCD (the PCD+ strain), while one does not (the PCD-strain).

The ABM progresses via discrete time steps, during which cells and patches update the state variables of itself and other agents in random order (see Supplementary material for computer code). Every time step there is a fixed probability that a disaster will occur and eliminate the susceptible phenotype. The choice of which phenotype is affected (what we referred to as A and B in the analytical model described in previous sections) is determined randomly and unbiasedly. Following the disaster, if there is space available in the patch, cells reproduce and give rise to an alternative phenotype with a constant probability. After reproduction, cells can migrate to new patches according to a fixed probability which is the same for all cells. At the end of a time step, cells of the PCD genotype undergo cell death probabilistically. Simulations end when a genotype goes extinct. We calculate *r* as the average probability a PCD+ cell that dies in a patch will be replaced by another PCD+ cell.

Disaster influences assortment by determining the relative benefit of phenotypic diversification through PCD. This follows from the analytical work and can be seen in the ABM simulations (Figures 5b and 5c). Overall, the relative performance of the PCD strain increases with the frequency of disasters. Not only do disasters select for rapid phenotypic diversification, but they also reduce the population to a relatively sparse state. Since the effects of birth on assortment/structure outweigh those of migration at low population densities, spatial structure (characterized by *r*) increases following a disaster. Thus, more frequent disasters increase the relative benefit of phenotypic diversity as well as the mean value of spatial structure (Figure 5d), both of which increase the benefit of PCD. The benefits of PCD are reduced in environments with very high disaster probabilities, as carrying capacity is rarely met so PCD offers fewer opportunities for increased reproduction, and more frequent disaster increases the stochasticity of the simulation. In between disasters, the PCD-strain increases in frequency and the value of *r* decreases. This happens because cells migrate which dissipates community structure, and also because PCD-cells occasionally replace dead PCD+ cells. This process would ultimately drive PCD+ strains extinct if it were not for disasters which favor phenotypic diversification (and thus PCD) and create spatial structure.

## Discussion

While recent years have seen a growing awareness that programmed cell death (PCD) is widespread among unicellular organisms [15, 16], few evolutionary hypotheses for its origin have been theoretically or experimentally investigated. Here we propose a novel hypothesis for the evolution of PCD in unicellular organisms: it increases the efficacy of microbial bet-hedging by creating generational turnover that yields increased phenotypic diversity. Using a simple bet-hedging model, we show analytically and computationally that PCD can be adaptive. In our model, PCD is advantageous because it allows microbes to become more phenotypically diversified, which is adaptive in the face of environmental uncertainty. It can be costly, however, if cells that die provide reproductive opportunities for competitors, not kin. In general, we find that fairly high rates of PCD can be favored by selection assuming low rates of stochastic phenotype switching (making population turnover from PCD more valuable) and high levels of genotypic assortment caused by spatial structure in the population (allowing the genotype undergoing PCD to capture most of the resources liberated by cellular suicide). Using an agent-based model we show that high rates of genotypic assortment for PCD strains (≈ .9) can emerge through normal demographic processes of reproduction and the bottlenecking effects of disasters.

**Figure 5:**
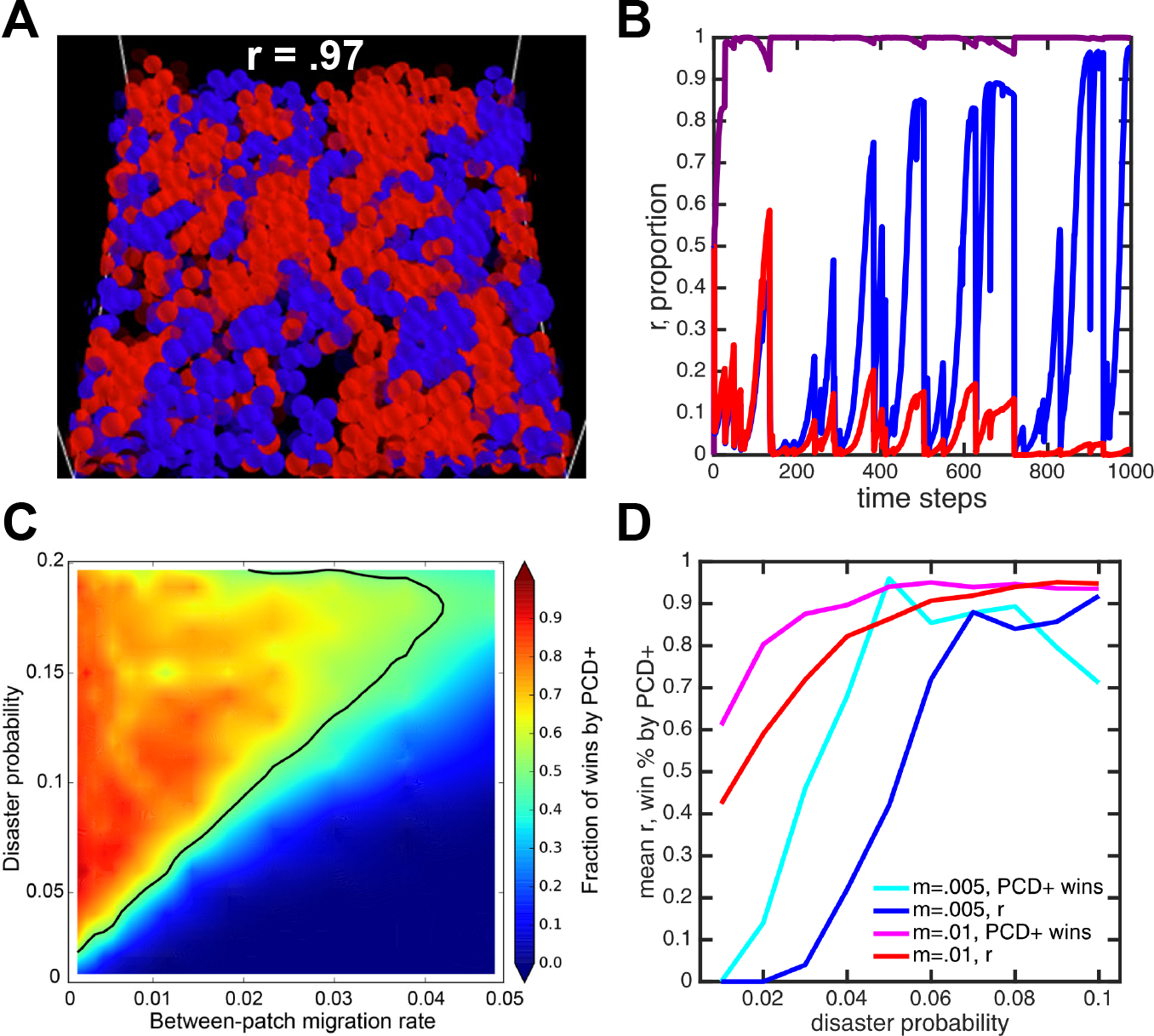
Generation of structure (*r*) via population processes. A) The 3D ABM shows that blue (PCD+) and red (PCD-) cells readily assort to create spatial structure. B) A characteristic time series of an ABM simulation showing the proportion of PCD+ (blue) and PCD-(red) as a function of the global carrying capacity as well as the value of *r*. Populations decrease following disasters. The ensuing recovery increases the value of *r*. Migration and PCD lead to a lowering of *r* and a decrease in the PCD+ strain. C) A contour plot shows the fraction of simulations won by PCD+ as a function of disaster probability and migration. More frequent disasters favor the PCD+ strain, but this is undermined by high rates of between-patch migration. The solid line represents equal competitiveness between PCD+ and – strains. 100 simulations were run for each of the 340 parameter combinations. D) The mean value of r and the fraction of wins by PCD+ are shown for two different migration probabilities. In both cases, *r* and the fraction of wins increases with disasters.

Due to the seemingly selfless nature of PCD, potential explanations have typically been sought at levels below (e.g. gene-[32] or phage-level; [33]) and above (e.g. colony, lineage-or population-level; [18, 22, 34]) that of the cell. It is also possible that the appearance of evolved PCD may occasionally arise as a byproduct or “malfunction” of traits that may confer cell-level advantages under certain environmental conditions [15]. Extant hypotheses for the adaptive significance of microbial PCD propose that it can provide novel functionality, coordinating “prokaryotic developmental programs” [34, 35], directing resources to mutant lineages that may be more capable of surviving a changing environment [36], or limitings the spread of pathogens to relatives [37]. Our hypothesis, that PCD can make bet-hedging strategies more effective, is both distinct from these alternatives, and may be fairly general, as stochastic phenotypic diversification is used to hedge against many different forms of risk [28, 38, 39, 40, 41, 42].

Why would organisms ever use PCD as a mechanism to diversify, when they could simply avoid the costs of cell death and evolve faster switch rates? We see two scenarios where PCD may be a useful tool. First, if evolving higher switch rates is either difficult or costly, PCD may offer a workaround. A number of cellular mechanisms allow for stochastic phenotype switching in microbes, including phase variation [43], contingency loci [44], and positive feedback loops in gene expression [45]. One of the elegant aspects of PCD is that it can increase the efficacy of any of these mechanisms without necessarily incurring complex pleiotropic side-effects. Second, PCD may allow for tuning of switch rates to match different environments that a microbe encounters. For example, if phenotypic diversification is beneficial only in challenging environments, but is otherwise costly, then PCD linked to an environmental cue correlated with the challenging environment may be a simple way to plastically increase diversification rates.

Diversification through PCD may be an alternative to anther common microbial bet-hedging mechanism: variable dormancy, in which only some cells of a clonal genotype become dormant [46, 47]. Individuals that become dormant, be it through persistence or the formation of resting stages like spores or cysts, gain resistance to a broad spectrum of environmental insults, but pay a considerable opportunity cost if the environment becomes favorable to growth [48]. Our model suggests that another, subtler price of dormancy is the missed opportunity to acquire resistance to future environmental stresses, through increased phenotypic diversification. Intriguingly, there is evidence that PCD delays the onset of dormancy (sporulation) during the transition to stationary phase in some bacteria by supplying resources to starving survivors [48]. Although PCD by a subpopulation may conceivably benefit surviving cells in the event of an abrupt shift back to favorable conditions [49], our model suggests that increased diversification may be another benefit of such population cycling.

Understanding the evolution of cellular suicide will require a plurality of approaches, informed by real-world ecology, backed by rigorous mathematical modeling and direct experimental investigation. We expect that there are many non-exclusive explanations for why microbes evolve PCD, but unfortunately few of the numerous hypotheses put forward in the literature have been mathematically investigated. Perhaps the main reason to model various hypotheses for PCD is to determine whether the conditions required for its evolution are permissive, or highly constrained. In this paper, for example, we find that high levels of PCD are only supported when the population is structured, but we also find that simple processes of death and clonal reproduction easily generate the necessary structure. While much work remains before we have a complete understanding of altruistic suicide, it is well worth the effort. Not only is it a topic of fundamental biological importance, it also has the potential to help generate novel therapeutic interventions [23, 50].

